# Luminance texture boundaries and luminance step boundaries are segmented using different mechanisms

**DOI:** 10.1101/2021.01.15.426873

**Authors:** Christopher DiMattina

## Abstract

In natural scenes, two adjacent surfaces may differ in mean luminance without any sharp change in luminance at their boundary, but rather due to different relative proportions of light and dark regions within each surface. We refer to such boundaries as *luminance texture boundaries* (LTBs), and in this study we investigate whether LTBs are segmented using different mechanisms than *luminance step boundaries* (LSBs). We develop a novel method to generate luminance texture boundaries from natural uniform textures, and using these natural LTB stimuli in a boundary segmentation task, we find that observers are much more sensitive to identical luminance differences which are defined by textures (LTBs) than by uniform luminance steps (LSBs), consistent with the possibility of different mechanisms. In a second and third set of experiments, we characterize observer performance segmenting natural LTBs in the presence of masking LSBs which observers are instructed to ignore. We show that although there may be some masking of LTB segmentation by LSBs, it is far less than that observed in a control experiment where both the masker and target are LSBs, and far less than that predicted by a model assuming identical mechanisms. Finally, we perform a fourth set of boundary segmentation experiments using artificial LTB stimuli comprised of differing proportions of white and black dots on opposite sides of the boundary. We find that these stimuli are also highly robust to masking by supra-threshold LSBs, consistent with our results using natural stimuli, and with our earlier studies using similar stimuli. Taken as a whole, these results suggest that the visual system contains mechanisms well suited to detecting surface boundaries that are robust to interference from luminance differences arising from luminance steps like those formed by cast shadows.

## INTRODUCTION

Understanding the mid-level visual representations which transform low-level features into high-level semantic representations of visual scenes remains an open question in vision science (**Pierce, 2015; Anderson 2020**). Over the years, multiple investigators have proposed that the purpose of mid-level vision is to build a representation of surfaces via a process of inference based on low-level feature detector outputs (**Marr, 1982; Nakayama, He, & Shimojo, 1995; Kubilius, Wagemans, & Op de Beeck, 2014; Anderson, 2020**). An essential operation for constructing surface-based representations is accurately detecting the boundaries separating different surfaces, an operation sometime referred to as *contour detection*, *image segmentation,* or *boundary segmentation* (**Malik, Belongie, Leung, & Shi, 2001; Arbelez, Maire, Fowlkes, Malik, 2010; Minaee et al., 2021; Zavitz & Baker, 2013, 2014; Pasupathy, 2015; DiMattina & Baker, 2019, 2021**). A number of local visual cues are potentially available at surface boundaries which can aid in boundary segmentation, including color, texture, and luminance differences (**Mely, Kim, McGill, Guo, & Serre, 2016; DiMattina, Fox, & Lewicki, 2012; Hansen & Gegenfurtner, 2009; Martin, Fowlkes, & Malik, 2004**), with luminance differences being one of the most reliable indicators of surface boundaries (**Mely, Kim, McGill, Guo, & Serre, 2016; DiMattina, Fox, & Lewicki, 2012**). However, in natural scenes luminance differences can arise from cues which do not correspond to surface boundaries, for instance, specular reflections, changes in surface orientation, or cast shadows (**Vilankar, Golden, Chandler, & Field, 2014**; **Casati & Cavanaugh, 2019**). Therefore, it is of great interest to better understand how the visual system may be able to infer the underlying cause of a local change in luminance.

Consider the boundary shown on the left of **Fig. 1a**, in which the luminance is constant within each surface, and there is a sharp change in luminance at the boundary. We refer to this stimulus as a *luminance step boundary* (LSB), and such a boundary can potentially arise from a variety of causes, including a boundary between two smooth surfaces, or a cast shadow falling on a single surface. However, for the boundary shown on the right of **Fig. 1a** arising from two juxtaposed natural surfaces, the luminance level varies greatly within each surface, as well as between the two surfaces, and there is no sharp change at the boundary. We refer to this second stimulus as a *luminance texture boundary* (LTB), since the luminance cue is defined by the different relative proportions of dark and light regions inside each texture, and is only clearly visible when integrating over a large spatial scale. **Fig. 1b** shows an example of a synthetic LTB used in previous work (**DiMattina & Baker, 2021**) in which the boundary visibility is determined by the relative proportions of white and black micro-patterns on opposite sides of a right-oblique boundary. Note that for these artificial stimuli that there are no texture cues apart from those defined by differences in luminance, thereby isolating this dimension of texture from other commonly studied dimensions like contrast (**DiMattina & Baker, 2019**), orientation (**Wolfson & Landy, 1998**) or density (**Zavitz & Baker, 2014**). This second kind of boundary (LTB) would typically only arise from a change in surfaces, or change in material properties, and would not be caused by things like cast shadows, changes in surface orientation with respect to the illuminant, or specular reflection. Therefore, any neural mechanism which responds differentially to the two kinds of luminance boundaries (LTB, LSB) would be potentially useful for distinguishing surface boundaries from cast shadows.

**Figure 1:**
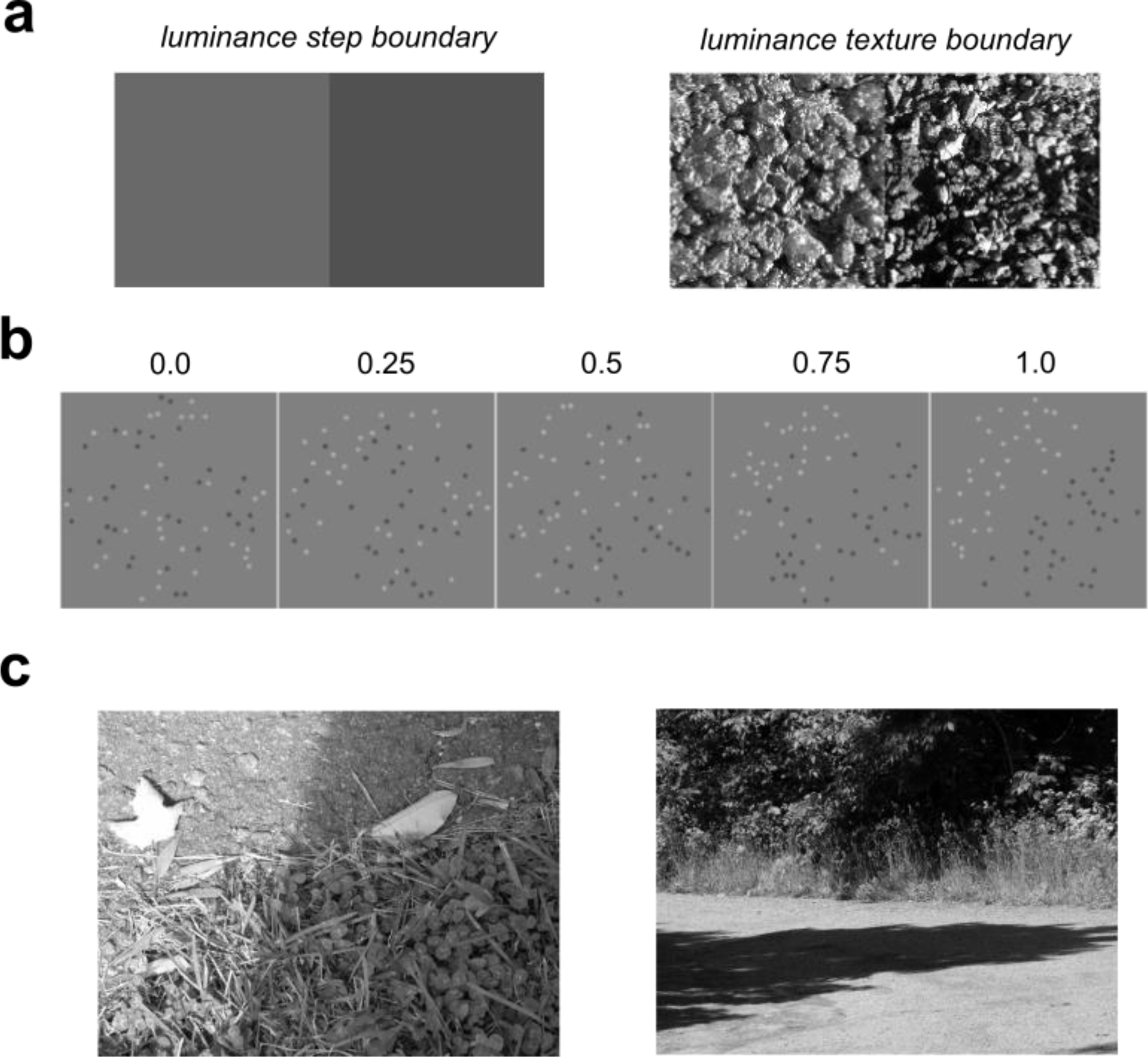
Examples of luminance step boundaries and luminance texture boundaries. (a) *Left:* A luminance step boundary (LSB). *Right:* A luminance texture boundary (LTB). Although the LTB exhibits a clear difference in average luminance between regions, there is substantial luminance variability within each region, with no sharp change in luminance at the boundary. (b) Artificial LTB stimuli formed by placing differing proportions of black and white micro- patterns on opposite sides of a disc. Boundary visibility is parameterized by the proportion π_*U*_ of micro-patterns on each side which are not balanced by a pattern of the same polarity on the opposite size. π_*U*_ = 0 corresponds to no boundary, π_*U*_ = 1 to opposite polarities on opposite sides. (c) LSBs can arise not only from boundaries between distinct surfaces, but also from changes in illumination caused by cast shadows.

Previous work in our laboratory has indeed suggested that different underlying mechanisms may sub-serve LTB and LSB segmentation (**DiMattina & Baker, 2021**). In this earlier study, we examined the ability of observers to segment LTBs in the presence of interference from masking LSB stimuli, and we found that we could best explain the results with a “Filter-Rectify-Filter” (FRF) model similar to those commonly utilized to model second-order vision (**Landy & Graham, 2004; Zavitz & Baker 2013, 2014; Victor, Conte, & Chubb, 2017; DiMattina & Baker, 2019**). By contrast, LSB segmentation was well explained with a simpler model positing a single stage of linear filtering implementing a simple luminance difference computation. However, in this earlier study of LTB segmentation, the masking LSB was only presented at threshold, whereas in natural vision one commonly segments surfaces in the presence of interference from clearly visible luminance steps created by cast shadows (**Casati & Cavanagh, 2019**), as shown in **Fig. 1c**. Another limitation in our previous work is that we only considered artificial luminance texture boundaries like those shown in **Fig. 1b**. Although such stimuli have the advantage of being easy to manipulate in a precise manner, it would also be desirable to see if our results generalize to more naturalistic stimuli formed by adding different kinds of luminance cues (either texture or steps) to uniform natural textures.

In the present study, we adduce further evidence that luminance steps and luminance textures are segmented via different mechanisms. In the first set of experiments (**Experiment 1**), we develop a novel method for creating luminance texture boundaries from uniform natural textures, and show that for these LTB stimuli, segmentation thresholds are significantly lower than the segmentation threshold for natural textures with super-imposed luminance step boundaries (LSBs). This difference in segmentation thresholds is consistent with the hypothesis that these two kinds of luminance boundaries are detected by different mechanisms. In a second and third set of experiments (**Experiments 2, 3**), we consider the segmentation of LTB stimuli in the presence of supra-threshold masking luminance step boundaries (LSBs) of the kind that might be caused by a cast shadow. We find that observers can quite successfully ignore a masking LSB when segmenting LTB targets, but not when segmenting LSB targets. Although we observed fairly weak masking of LTB targets by LSB maskers, we found that these effects (when present) are dependent on the relative orientations and phases of each kind of boundary, suggesting some degree of interaction between the two cues. Finally, in a fourth set of experiments (**Experiment 4**), we repeat the second set of experiments using the artificial luminance texture boundaries investigated previously (**DiMattina & Baker, 2021**), obtaining similar results.

Taken as a whole, the present experiments utilizing natural stimuli point to the same conclusion as those with artificial stimuli, namely that different underlying mechanisms are responsible for representing different kinds of luminance boundaries (**DiMattina & Baker, 2021**). Since luminance step boundaries often arise from cast shadows rather than surface boundaries (**Fig. 1c**), the existence of mechanisms which are sensitive to luminance texture differences but not luminance steps may serve an important functional role for segmenting natural visual scenes.

## METHODS

### Stimuli and Task

#### Natural luminance texture boundary stimuli

Pairs of natural textures will generally differ from each other in a variety of dimensions unrelated to luminance differences (**Fig. 1a**, *right*). Therefore, in order to isolate luminance cues from other textural cues (e.g., orientation, density, spatial frequency), we developed a novel procedure to synthesize luminance-defined texture boundaries from uniform natural textures. The basic idea is that luminance-defined texture differences can be created by increasing the brightness of the “bright” areas on one side of the boundary (leaving the dark areas on this side the same), and increasing the darkness of the of “dark” areas on the other side of the boundary (leaving the light areas the same). This is by no means the only way to create a texture-defined luminance difference, but it roughly models the common situation where one surface has more “bright” areas and fewer “dark” areas than an adjacent surface (e.g., **Fig. 1a**, *right*).

Textures were obtained from the Brodatz database (**Brodatz, 1966**). The particular textures we used in the present study were from the Normalized Brodatz Texture (NBT) Database (https://multibandtexture.recherche.usherbrooke.ca/normalized_brodatz.html), a modification of the original Brodatz database where each texture was transformed to occupy the entire 256 level grayscale range fairly uniformly, making it impossible to discriminate between these textures using only first-order statistics. From the full set of textures, we selected a subset of N = 7 textures for this study, shown in **Fig. 2c**. These textures were chosen because they were relatively uniform, non-oriented, and had small micro-patterns relative to the texture size. Sampled texture patches were 96x96 sub-images taken from random locations in the 128x128 texture images. From the original texture patch, a “left” and “right” side image (both centered at zero) was created by multiplying the original image by an obliquely oriented (left or right-oblique), cosine-tapered modulation envelope which goes zero on the opposite size of the boundary, such that adding up the two sides yielded the original image (**Fig. 2a**). A desired luminance difference across the boundary (ΔL, dimensionless units) was created by centering the stimulus at zero mean, and then scaling the positive pixels on one side of the boundary, and the negative pixels on the opposite side, by a factor *k* > 0 obtained via numerical optimization in order to set ΔL to some desired value. For each stimulus, we also specified a phase parameter, which specified whether the left (phase = 0 deg.) or right (phase = 180 deg.) side was brighter. This synthesis procedure is illustrated schematically in **Fig. 2a** for Brodatz texture **D111**. The final stimulus was then set to mean luminance of 0.5 (screen mid-point) and scaled to 256 x 256 to subtend 4 dva on our display. The bottom row of **Fig. 2b** shows LTBs synthesized from Brodatz texture **D28** using our procedure at four different levels of luminance difference (ΔL = 0, 500, 1000, 1500).

**Figure 2:**
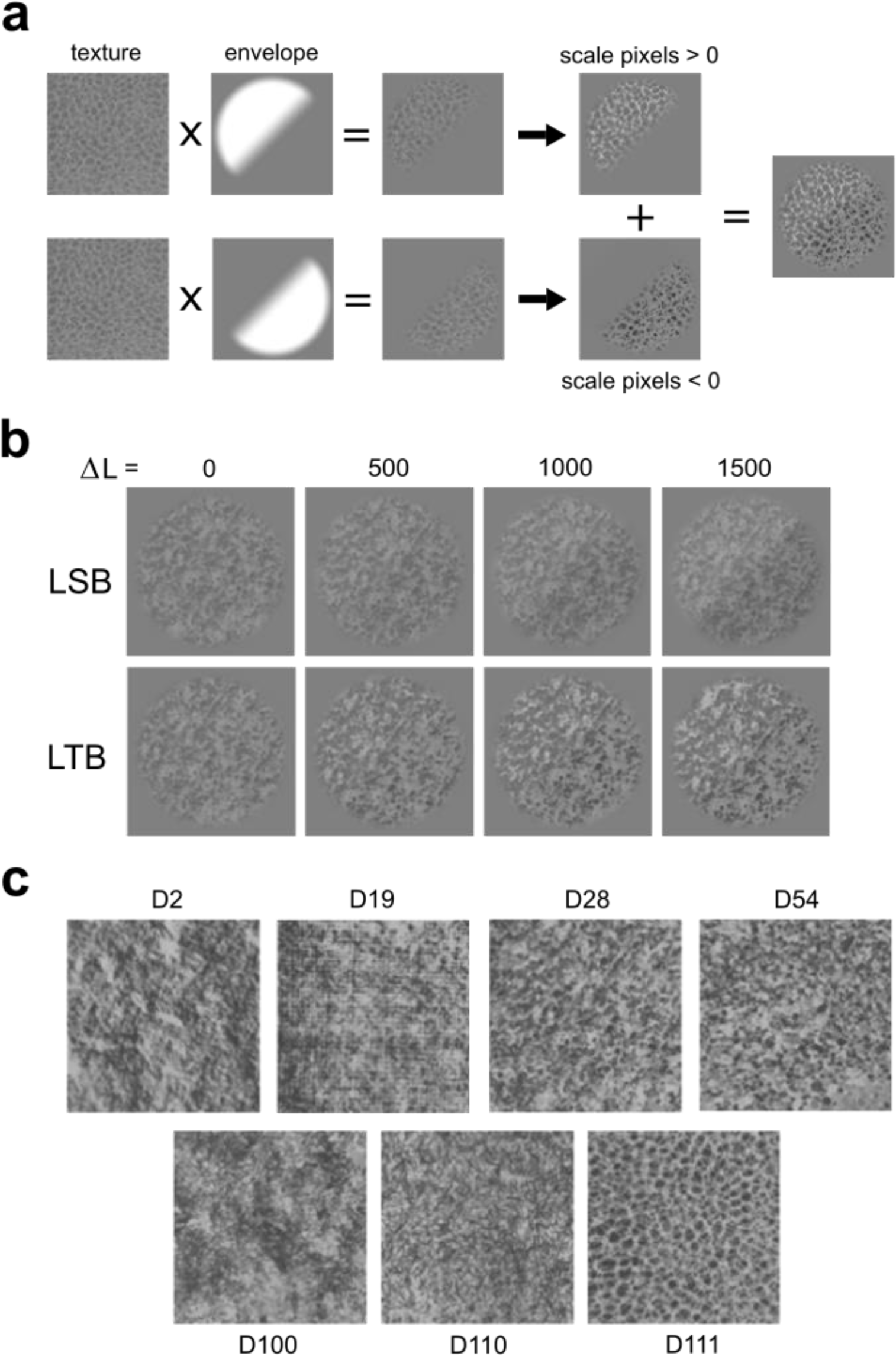
Natural luminance texture boundary (LTB) synthesis. (a) Illustration of our synthesis procedure for natural LTB stimuli. A zero-mean uniform texture (*left column*) was first multiplied by an envelope (*center-left*) to create two halves (*center*). In one half, the pixels greater than zero were scaled, while those less than zero were left unchanged. In the other half pixels less than zero were scaled, while those greater than zero were left unchanged (*center-right*). The two scaled stimuli were then added together to create a luminance boundary (*right*). (b) Texture **D28** with a luminance step boundary added (*top row*), and with a luminance texture boundary generated using our procedure (*bottom row*). In each row, the luminance difference across the diagonal (dimensionless units ΔL) goes from 0 to 1500 in steps of 500 (ΔL = 0:500:1500). (c) The full set of N = 7 Brodatz textures used in this study.

#### Artificial luminance texture boundary stimuli

Artificial luminance texture boundary (LTB) stimuli (**Fig. 1b**) were created by placing different proportions of black (B) and white (W) micro-patterns on opposite halves of a circular disc. The visibility of the boundary was determined by the relative proportions (π_*U*_) of white and black micro-patterns on each side of the boundary which are *not* balanced by a micro-pattern on the opposite side having the same polarity. The proportion of unbalanced patterns π_*U*_ can range from 0 (no boundary) to 1 (opposite colors on opposite sides). These stimuli have been employed in previous work (**DiMattina & Baker, 2021**), and examples with varying levels of π_*U*_ are shown in Fig. 1b.

The artificial LTB stimuli had either 32 or 64 eight-pixel Gaussian (σ = 2 pixels) micro- patterns on each side of the boundary. Maximum micro-pattern amplitude *A* was set to +/- 0.15 (W/B) dimensionless units with respect to the neutral gray mid-point (0.5), with Michelson contrast given by *c*_*M*_ = 2*A*. For this level of *A*, the micro-patterns were clearly visible. By design, the stimuli had no luminance difference across the diagonal perpendicular to the boundary (which we refer to as the *anti-diagonal*), and stimulus phase was set to either 0 or 180 degrees (left/right side brighter). Micro-pattern locations were chosen pseudo-randomly, with overlap being prevented by maintaining a list of locations which are currently unoccupied and hence available to put new micro-patterns. Artificial LTB stimuli were also 256x256 and subtended 4 deg. of visual angle.

#### Luminance step boundaries

Luminance step boundaries (LSB) like those shown in **Fig. 3a** were created by multiplying an obliquely oriented step edge by a cosine-tapered circular disc. To better model the penumbral blur characteristic of shadows (**Casati & Cavanagh, 2019**), the luminance level transitioned from low to high (cosine envelope) over the central 20% of the boundary. The detectability of this edge was varied by manipulating its Michelson contrast (*c*_*M*_), and again the envelope phase was randomized.

**Figure 3:**
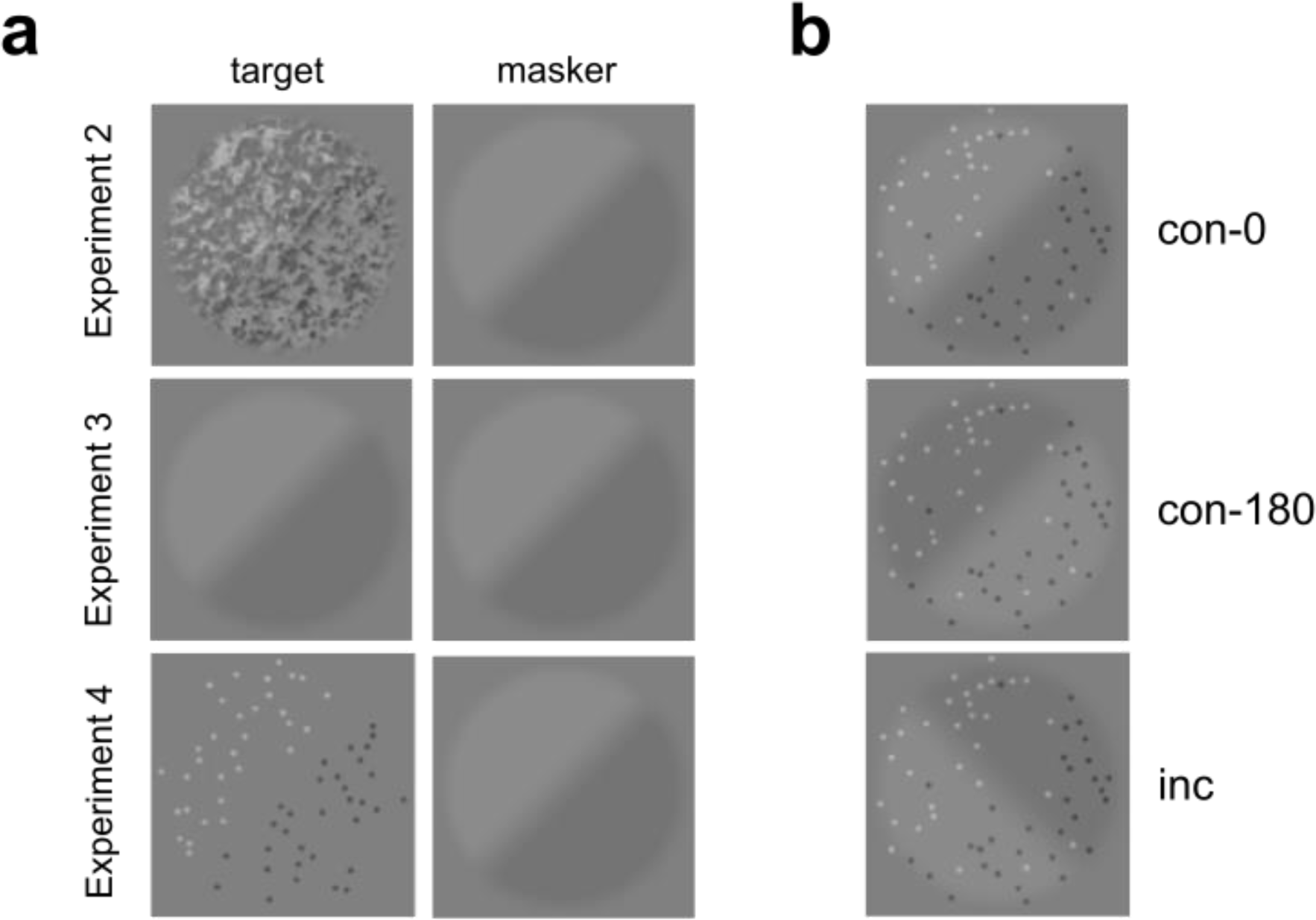
Nature of the target and masking stimuli for **Experiments 2-4**. (a) In **Experiment 2** (*top*) and **Experiment 4** (*bottom*), observers indicated the orientation (left/right-oblique) of a natural or artificial LTB while ignoring a masking LSB. In a control experiment (**Experiment 3**), both target and masker were LSBs. (b) Illustrations of phase relations between a target LTB (dots) and masking LSB. In the incongruent (**inc**) case (bottom), masker and target indicate opposite boundary orientations. In the congruent (**con**) case where the boundary orientation is the same, it is either phase-aligned (**con- 0,** top) or opposite phase (**con-180,** middle).

In some experiments, the LSB was presented in isolation, and in others it was added to a uniform texture (**Fig. 2b**, *top*), with its amplitude numerically optimized to give rise to a specified luminance difference across the diagonal (ΔL). LSB stimuli were also 256 x 256 pixels and scaled to subtend 4 deg. of visual angle.

#### Psychophysical task

Observers performed a single-interval segmentation task identifying the orientation of a boundary (LTB or LSB) as either right-oblique (+45 deg. with respect to vertical, **Fig. 1b**) or left-oblique (- 45 deg. w.r.t. vertical). This *orientation-identification* task has been used in a number of previous studies investigating boundary segmentation **(Arsenault, Yoonessi, & Baker, 2011**; **Zavitz & Baker, 2013, 2014; DiMattina & Baker, 2019, 2021**). Boundary visibility was adjusted for each type of stimulus (LSB, LTB) using a 1-up, 2-down staircase procedure, which focuses trials at stimulus levels leading to 70.71% correct performance (**Leek, 2001**). Stimulus levels were chosen for all experiments so that observer performance varied smoothly from chance to perfect or nearly perfect, which ensured robust and accurate estimates of thresholds with our staircase. For the artificial stimuli, boundary visibility can be varied using definitional units (LTB: π_*U*_, LSB: *c*_*M*_).

However, for the natural LTB stimuli, boundary strengths can only be specified in units of (dimensionless) luminance difference ΔL across the diagonal.

### Observers

Author CJD and two naïve undergraduate student researchers (MXD, BNG) served as observers in these experiments. All observers had normal or corrected-to-normal visual acuity. All observers gave informed consent, and all experimental procedures were approved by the FGCU IRB (Protocol 2014-01), in accordance with the Declaration of Helsinki.

### Visual Displays

Stimuli were presented in a dark room on a 1920x1080, 120 Hz gamma-corrected Display++ LCD Monitor (Cambridge Research Systems LTD®) with mid-point luminance of 100 cd/m^2^. This monitor was driven by an NVIDA GeForce® GTX-645 graphics card, and experiments were controlled by a Dell Optiplex® 9020 running custom-authored software written in MATLAB® making use of Psychtoolbox-3 routines (**Brainard, 1997; Pelli, 1997; Kleiner et al., 2007**). Observers were situated 133 cm from the monitor using a HeadSpot® chin-rest, so that the 256x256 stimuli subtended approximately 4 deg. of visual angle.

## Experimental Protocols

### Experiment 1: Comparing LTB and LSB and segmentation thresholds

In the first set of experiments, we compared the segmentation thresholds (in units of ΔL) for two different kinds of luminance boundaries created from natural textures: Luminance texture boundaries (**Experiment 1a**), and luminance step boundaries created by adding uniform luminance steps to natural textures (**Experiment 1b**). Examples of both kinds of boundary (LTB, LSB) are shown in **Fig. 2b**.

When working with uniform natural textures, although we specify the luminance across the diagonal, because of random variability in the texture there will generally also be a very small luminance difference across the *anti-diagonal* (the boundary orthogonal to the direction of luminance modulation). Although this minor variation will not give rise to any kind of visible boundary, in order to ensure that there was zero luminance difference cue across the anti-diagonal, for **Experiments 1a** and **1b** we added a small luminance step (LSB) across the anti-diagonal to nullify regional luminance differences. We refer to these stimuli as *balanced* (**bal**). As we can see from the stimuli in **Fig. 2b**, all of which are balanced, this added LSB does not give rise to any visible boundary across the anti-diagonal. Nevertheless, as a control we also repeated **Experiments 1a, b** without nullifying the small random luminance difference across the anti-diagonal (**Experiments 1c, d**), a condition we call *unbalanced* or *raw* (**raw**). As detailed in the Results, no difference was observed between these conditions.

In all conditions of **Experiment 1**, observers CJD and MXD were tested using the staircase procedure defined on a set of at N = 15 luminance difference levels ΔL from 0 to 1400 (dimensionless units) in steps of 100 (which we denote ΔL = 0:100:1400). Observer BNG was tested using a broader range of N = 21 levels (ΔL = 0:100:2000).

### Experiment 2: Masking Natural LTBs with LSBs

If luminance texture boundaries are detected using different mechanisms than luminance step boundaries, we should expect that observers should be able to segment luminance textures while ignoring super-imposed luminance step boundaries. In a previous study aimed at testing competing hypotheses of LTB segmentation mechanisms (**DiMattina & Baker, 2021**), we demonstrated this to be the case for artificial luminance texture boundaries in the presence of luminance step boundaries presented at threshold. However, in that study, we did not consider luminance step boundaries presented above their segmentation threshold. Since cast shadows give rise to supra- threshold LSBs which do not correspond to surface boundaries (**Fig. 1c**), it is of interest to see if LTB segmentation is robust to interference from supra-threshold LSBs. Likewise, in this previous study we only made use of artificial luminance texture boundary stimuli (**Fig. 1b**) as opposed to natural luminance texture boundaries (**Fig. 2**).

In **Experiment 2a**, three observers segmented a subset of the natural LTBs used in **Experiment 1** (**D2, D28**, **D54**), but this time in the presence of supra-threshold masking LSBs presented at 2, 4 and 6 times the JND for segmenting the LSB in isolation. Observers were instructed to ignore the LSB and segment the LTB (indicate its orientation as L/R-oblique). This paradigm is illustrated in **Fig. 3a** (*top*). Each observer performed 600 trials using a staircase procedure defined on the levels ΔL = 0:100:1800. Of these 600 trials, for 300 trials the masking LSB had the same orientation as the target LTB, and we refer to these trials as *congruent* (**con**). For the remaining 300 trials, the masking LSB had the opposite orientation as the target LTB, and we refer to these trials as being *incongruent* (**inc**). Within the set of 300 congruent (**con**) trials, for 150 trials the masking LSB had the same phase (**con-0**) as the target LTB, and for 150 trials the masking LSB had an opposite phase (**con-180**). These conditions (**inc**, **con-0**, **con-180**) are all illustrated (for an artificial LTB) in **Fig 3b**.

In **Experiment 2a**, we balanced stimuli so that there was no luminance difference across the anti-diagonal (**bal**), as **Experiment 1** suggested that this had no influence on LTB segmentation performance. However, for one of the three textures (**D54**), we ran an additional control condition (**Experiment 2b**) in which we did not apply our balancing procedure (**raw**).

### Experiment 3: Masking LSBs with LSBs

In another control experiment (**Experiment 3)** illustrated in **Fig. 3a**, observers segmented a target LSB in the presence of a masking LSB presented at the same levels as in **Experiment 2**. This experiment is illustrated schematically in **Fig. 3a** (*middle*). The purpose of this experiment was to provide a baseline for interpreting the results of **Experiment 2**, since in **Experiment 3** both the masker and target are of the same kind and will necessarily engage identical neural mechanisms. If degree of masking is comparable between **Experiments 2** and **3**, this suggests that the same mechanisms are used for both LTBs and LSBs. By contrast, if we observe much less masking in **Experiment 2** than **Experiment 3**, it suggests different mechanisms for the LTB stimuli. Observers MXD and BNG performed 600 trials with a staircase procedure defined on on stimulus levels ΔL = 0:100:2600. Observer CJD performed 400 trials on levels ΔL = 0:100:1800.

### Experiment 4: Making artificial LTBs with LSBs

**Experiment 4** was identical to **Experiment 2**, except in this case the luminance texture boundaries were artificial LTBs (**Fig. 1b**) used in our previous studies (**DiMattina & Baker, 2021**). By design, these stimuli only had luminance differences across the diagonal, with zero luminance difference across the anti-diagonal. Artificial LTBs were tested at two micro-pattern densities: *n*_*p*_ = 32 and *n*_*p*_ = 64 gaussian “dot” micropatterns per side. Boundary visibility was changed by setting the value of the parameter π_*U*_ (proportion unbalanced micropatterns), which was varied from π_*U*_ = 0: 0.125: 1 (*n*_*p*_ = 32) or π_*U*_ = 0: 0.0625: 1 (*n*_*p*_ = 64). Each observer performed 600 trials, with the relative orientations and phases of the LTB target and LSB masker (presented at 2, 4, 6 JND) being the same as in **Experiment 2**. This experiment is illustrated schematically in **Fig. 3a** (*bottom*).

### Data Analysis

#### Psychometric function fitting and bootstrapping

To determine thresholds (75% performance) in each experimental condition, data was fit using a sigmoid psychometric function (PF) using MATLAB routines from the Palemedes Toolbox (http://www.palamedestoolbox.org/), described in **Prins & Kingdom (2018)**. The proportion correct responses (*p*_*c*_) to a stimulus of intensity *x* is given by:

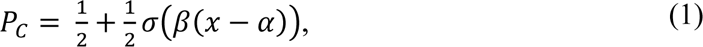

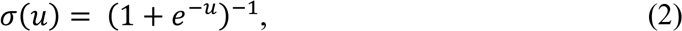

where β is the slope, and α the threshold.

Bootstrapping (**Efron & Tibshirani, 1994**) was employed to plot non-parametric 68% and/or 95% confidence intervals for the psychometric function thresholds. For bootstrapping analyses, N = 2000 simulated datasets were created as follows: For each stimulus level with *ni* presentations and *ci* experimentally observed correct responses (proportion of correct responses *pi* = *ci*/*ni*), we sampled from a binomial distribution having *ni* trials with probability *pi* to create a simulated number of correct responses for that stimulus level. We fit our model (1) to each of these simulated datasets, and obtained a distribution of estimates of the stimulus levels corresponding to JND (75% correct) performance. After a conservative outlier removal procedure (removing observations 3 IQR away from the median), we computed the median of the final distribution as well as the ranges of scores containing 68% and 95% of the data. We refer to these ranges as bootstrapped confidence intervals.

### Generic masking model

In order to compare the observed threshold elevations to the theoretical elevations we would expect when the masker and target are processed by identical mechanisms, we derived the following simple model. We assume that the observer computes a decision variable *u* = *u*_*R*_ − *u*_*L*_ by taking the difference of two mechanisms, one of which is sensitive to a right-oblique (R) boundary and the other a left-oblique (L) boundary. We further assume that each mechanism produces a noisy Gaussian output with unit variance and stimulus-dependent means. The final decision variable *u* is therefore Gaussian distributed with mean *μ*_*R*_ − *μ*_*L*_ and variance *σ*^2^ = 2. Assume without loss of generality that the target is right-oblique, and the mean response of the R mechanism is given by a power-law transducer function *μ*_*R*_ = [*gx*]^α^ , where *x* denotes the stimulus intensity, *g* the gain and *α* the exponent. In the absence of a mask *μ*_*L*_ = 0, and the probability *p*_*c*_ of making a correct response is *p*(*u* > 0), given by the expression:

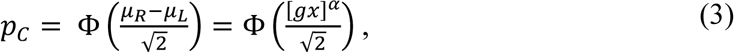

and JND performance level *J* (typically *J* = 0.75) is obtained at level 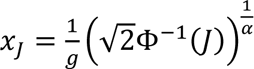. Now, assume we have a left-oblique mask set at level *x*_*M*_, so that *μ*_*L*_ = [*gx*_*M*_]^α^. For a right-oblique target, the mean output of the right-oblique mechanism is given by *μ*_*R*_ = [*gx*]^α^, and from (3) we obtain JND performance level *J* for

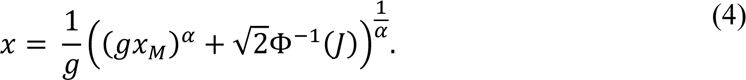

Following others, we assume fixed internal noise for each mechanism which is independent of stimulus level (e.g., **Legge & Foley, 1980**). Although this assumption has been somewhat a point of debate (**Georgeson & Meese, 2006**), recent work re-analyzing scaling and discrimination data (**Whittle, 1986, 1992**) has provided strong evidence that internal noise is independent of stimulus level (**Kingdom, 2016**).

In our analyses, we found α = 1 to provide an excellent fit to the data where the target and masker are both of the same kind (LSB) and therefore activate the same mechanisms (**Experiment 3**). This may seem somewhat surprising, since it is well-known that the transducer function for contrast discrimination is a near-miss to Weber’s law with exponent 0.7 (**Legge, 1981**). However, this law only holds for a function that is defined over the full range of contrast from 0 to 100%. Our LSB masking stimuli only covered a very narrow range of contrasts (1-6%), and over this highly restricted range the underlying transducer function is approximately linear (α= 1).

## RESULTS

### Experiment 1: Segmentation thresholds for natural LTB and LSB stimuli

In previous work using artificial stimuli like those in **Fig. 1b** (**DiMattina & Baker, 2021**), we obtained evidence consistent with the hypothesis that different neural mechanisms may be employed to segment luminance texture boundaries (LTBs) and luminance step boundaries (LSBs). In order to investigate this idea further using complex naturalistic stimuli, we developed a method for synthesizing LTBs from uniform natural textures, as illustrated in **Fig. 2a**. We compared the segmentation thresholds for these naturalistic LTBs (**Fig. 2b**, *bottom*) to those obtained from LSBs super-imposed onto textures (**Fig. 2b**, *top*), where in both cases we were able to precisely control the luminance difference ΔL across the diagonal. If there is only a single mechanism which computes luminance differences for segmentation, then we should expect that identical luminance differences ΔL should give rise to identical segmentation performance for LTB and LSB stimuli. By contrast, different thresholds for these two kinds of stimuli would be suggestive of different mechanisms being employed for each kind of boundary.

**Figs. 4, 5** summarize our results from **Experiment 1**. **Fig. 4a** demonstrates that the ΔL segmentation thresholds (75% JND) for an LSB super-imposed on a texture (blue: **bal**, green: **raw**) are significantly higher (lines show bootstrapped 95% confidence intervals) than thresholds for an LSB in isolation (black dashed lines). Plotting the elevation in both natural units (luminance difference ΔL) and JND units reveals that over 42 cases defined by the 3 observers, 7 textures, and 2 conditions (**bal**, **raw**), in every case but one (MXD, texture D111, **bal**), or 41/42 cases, we see a statistically significant threshold elevation (as determined by intersection with the 95% CI), with substantial variability from texture to texture for each observer (**Fig. 4a**). Comparing the difference in thresholds between the anti-diagonal balanced (**bal**) and anti-diagonal unbalanced (**raw**) cases reveals that over 3 observers and 7 textures, in the vast majority of cases (15/21) we do not see any significant difference (meaning zero lies outside the 95% confidence interval of the difference) between the thresholds obtained in the two conditions (**Fig. 4b**), with observer CJD accounting for 4 of the 6 significant differences.

**Figure 4:**
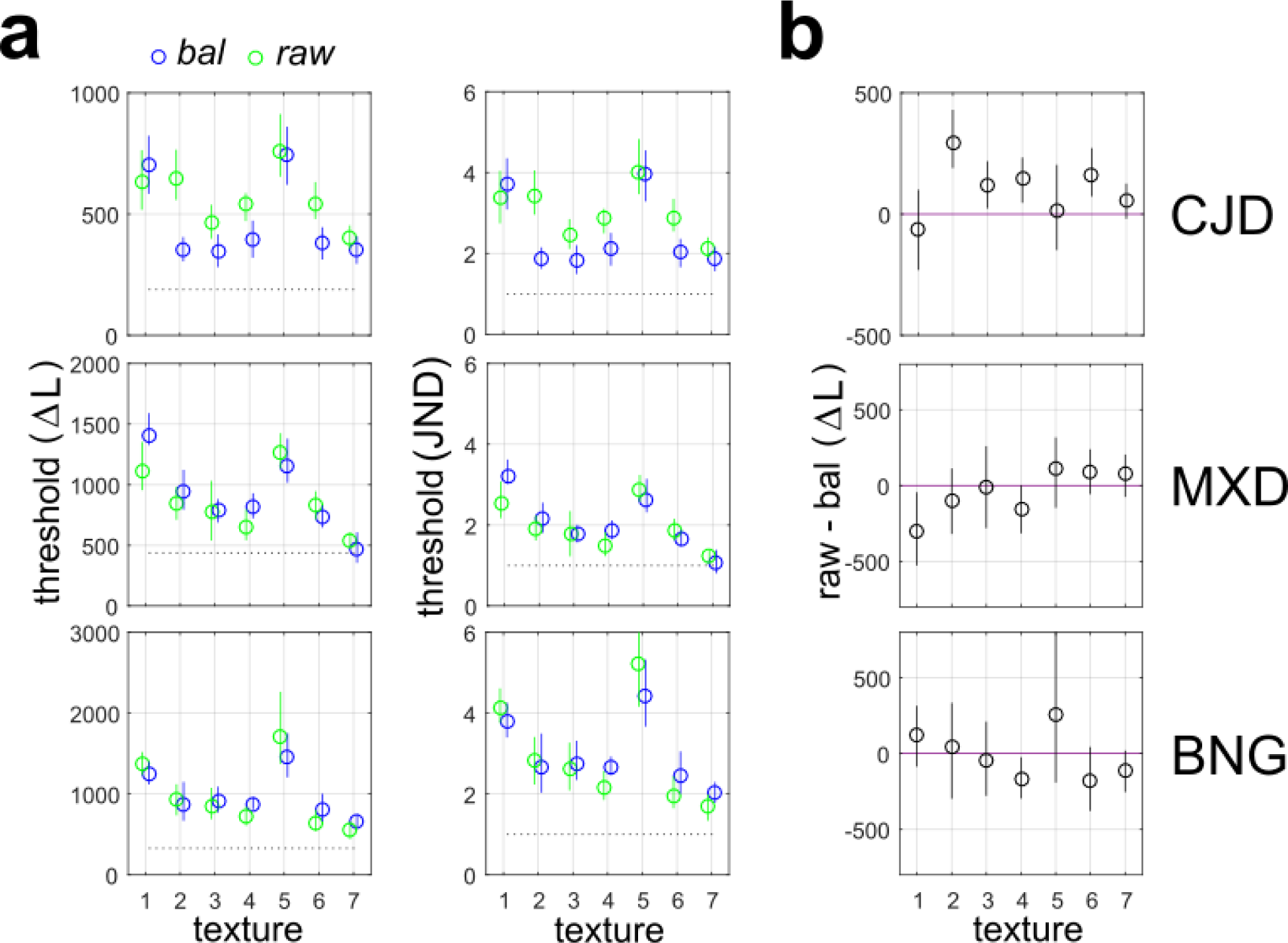
Segmentation thresholds for luminance step boundaries (LSBs). (a) *Left:* Thresholds for LSB super-imposed on each of the Brodatz textures in units of absolute luminance difference ΔL across the diagonal. Blue symbols indicate stimuli which are luminance- balanced across the anti-diagonal (**bal**), green symbols indicate stimuli which are not luminance- balanced across the diagonal (**raw**). The segmentation thresholds for LTBs in isolation (i.e., not super-imposed on a texture) are shown by dashed black lines. Symbols indicate medians, and error bars indicate bootstrapped 95% confidence intervals (N = 2000 bootstraps). *Right:* Same as left panels, but in units of the JND for the LTB presented in isolation. (b) Median differences between the raw and balanced thresholds in the segmentation task in units of ΔL. Error bars indicate 95% bootstrapped confidence intervals of the difference.

**Figure 5:**
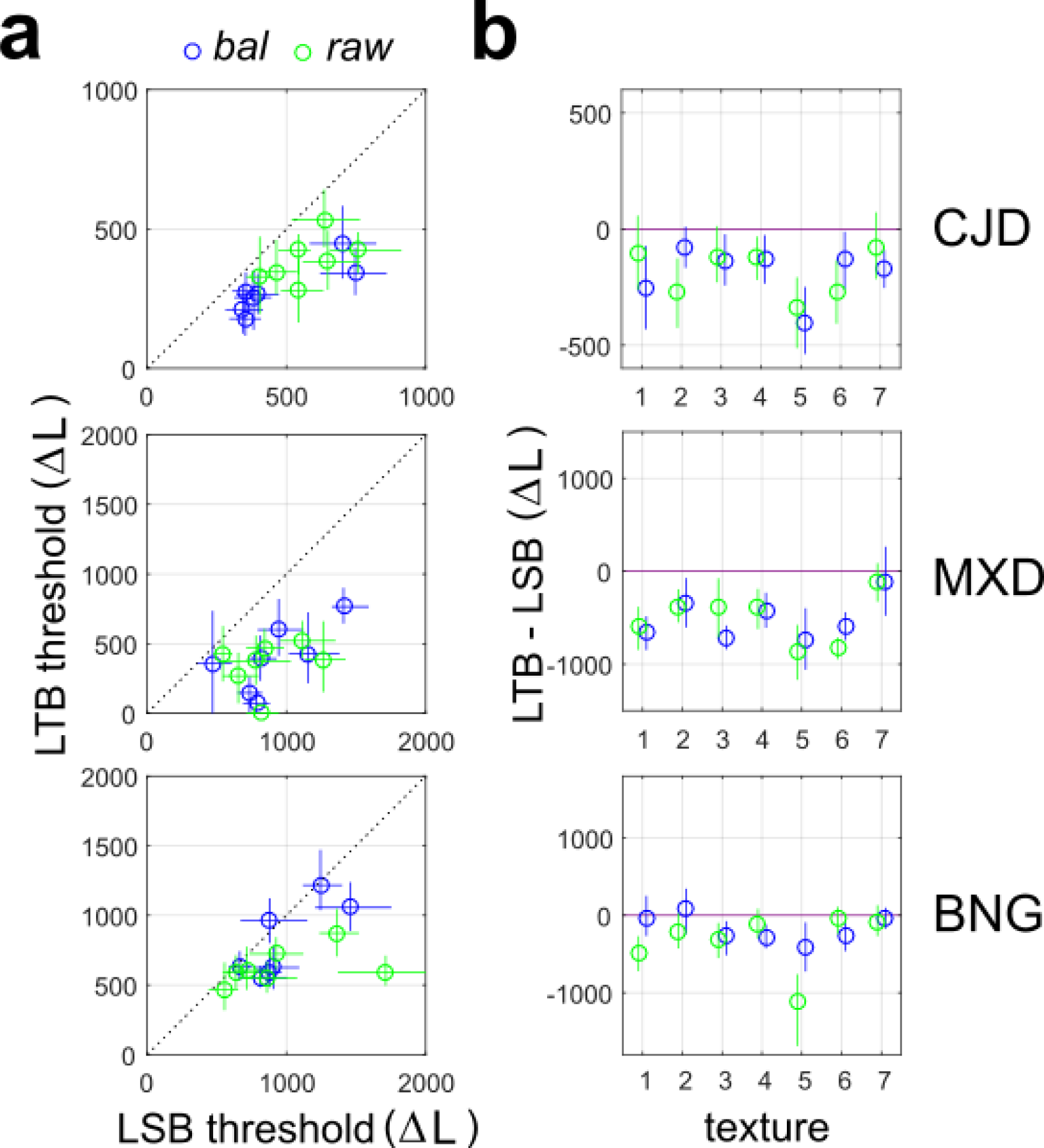
Segmentation thresholds for LTBs and LSBs. (a) Scatterplot of segmentation thresholds for LSB stimuli superimposed on natural textures and LTB stimuli generated from natural textures in units of luminance differences ΔL across the diagonal. Blue symbols indicate balanced stimuli (**bal**) modified to have zero luminance difference across the anti-diagonal, and green symbols indicate unbalanced stimuli (**raw**). Symbols indicate medians, and lines indicate 95% bootstrapped confidence intervals (N = 2000 bootstraps). (b) Median differences between thresholds for LSB and LTB stimuli for balanced (**bal**, blue) and unbalanced (**raw**, green). Lines indicate 95% bootstrapped confidence intervals of the difference.

Analyzing the median LSB segmentation thresholds for each of the N = 7 textures in the **bal** and **raw** conditions revealed significant differences from each subject’s LSB segmentation threshold in isolation for all observers (rank-sum test, N = 7, *p* = 0.016 for all observers/conditions). Comparing the LSB segmentation thresholds in the **bal** and **raw** conditions failed to yield a significant difference for any observer (sign-rank test, N = 7. CJD: *p* = 0.109, MXD: *p* = 0.578, BNG: *p* = 0.938).

**Fig. 5a** demonstrates that segmentation thresholds across textures and observers were often significantly lower for LTB stimuli than LSB stimuli, as we see all of the symbols falling either on or below the diagonal on a plot with the LSB segmentation threshold on the ordinate and LTB segmentation threshold on the vertical axis. This effect is consistent for both the **bal** (blue symbols) and **raw** (green symbols) conditions, with the lines denoting 95% confidence intervals for the threshold. **Fig. 5b** shows the difference between thresholds for the LTB and LSB stimuli in both the anti-diagonal balanced (**bal**) and unbalanced (**raw**) conditions. For observer CJD, the LTB segmentation threshold was significantly lower than the LSB threshold for 6/7 textures for the **bal** stimuli (blue symbols) and 4/7 for the **raw** stimuli (green symbols). For observer MXD, LTB threshold was lower than LSB for 6/7 (**bal**) and 6/7 (**raw**) textures, and for BNG, LTB thresholds were lower than LSB for 4/7 (**bal**) and 3/7 (**raw**) textures. Critically, in none of the 42 cases (3 observers, 7 textures, 2 balancing conditions) did the LSB segmentation threshold significantly exceed that of the LTB for the same texture (**Fig. 5b**). Comparing the median segmentation thresholds in for LTB and LSB stimuli across all N = 7 textures revealed statistically significant differences for observers CJD and MXD in the both the **bal** and **raw** conditions (sign-rank test, N = 7, *p* = 0.016, both observers and tests). For observer BNG, a significant difference was obtained in the **raw** (*p* = 0.016) but not **bal** (*p* = 0.078) condition.

Taking the results of **Experiment 1** as a whole, we conclude that observers are generally more sensitive to identical luminance differences across the diagonal when they arise from different relative proportions or local intensities of dark and light areas on each side of the boundary than when they arise from the addition of a uniform luminance step, as might arise from a cast shadow. This speaks to the possibility that different mechanisms may be employed for these segmentation tasks.

### Experiments 2, 3: Masking Natural LTBs with LSBs

Masking experiments have long been used in vision science to investigate whether or not two stimuli are processed using the same or different mechanisms. Perhaps the best example of this comes from classical work in spatial vision, which used masking paradigms (**Stromeyer & Julesz, 1972; Legge & Foley, 1981; DeValois & Switkes, 1983; Wilson, McFarlane & Phillips, 1983**) to help establish that the early visual system decomposes images into multiple spatial frequency and orientation channels using operations resembling a localized Fourier transform (**DeValois & DeValois, 1988**). Inspired by our results in **Experiment 1**, we next performed a masking experiment to further investigate the hypothesis of different mechanisms for LTB and LSB segmentation (**Experiment 2**).

In **Experiment 2a**, observers were instructed to segment LTBs in the presence of masking LSBs which they were instructed to ignore, as illustrated in **Fig. 3a**. In contrast to our previous study (**DiMattina & Baker, 2021**), here the masking LSBs were presented well above the segmentation threshold for the LSB in isolation (2, 4, 6 x JND). This paradigm models the situation where one is segmenting the boundary between two surfaces in the presence of a clearly visible cast shadow. In order to put the results of **Experiment 2a** in context, we performed a control condition (**Experiment 3**) in which observers segmented luminance steps (LSBs) in the presence of masking LSBs. Since both the target and the masker are of the same kind, by necessity they will activate the same mechanisms, and therefore this experiment provides us with an upper limit to the extent of possible masking.

**Fig. 6** shows the results of **Experiment 2** for each observer-texture pair. Here we plot the level of the LSB masker and LTB segmentation threshold in units of absolute luminance difference across the diagonal (ΔL). We see that the LSB does mask the LTB target to a moderate extent, as the target segmentation thresholds (blue symbols) are generally higher than thresholds in the absence of an LSB masker (blue dotted line, **Experiment 1**), and increase with increasing masker levels. For observer CJD, over 3 textures and 3 masker levels, in 8/9 cases there is statistically significant elevation of the segmentation threshold (i.e., the threshold in the absence of a masker does not lie in the 95% CI). For observer BNG, we see 7/9 cases with significant elevation. MXD was a little more robust to LSB masking (4/9 cases), and for one texture (**D54**) MXD exhibited no significant increases in threshold relative to the unmasked condition for any tested masker level.

**Figure 6:**
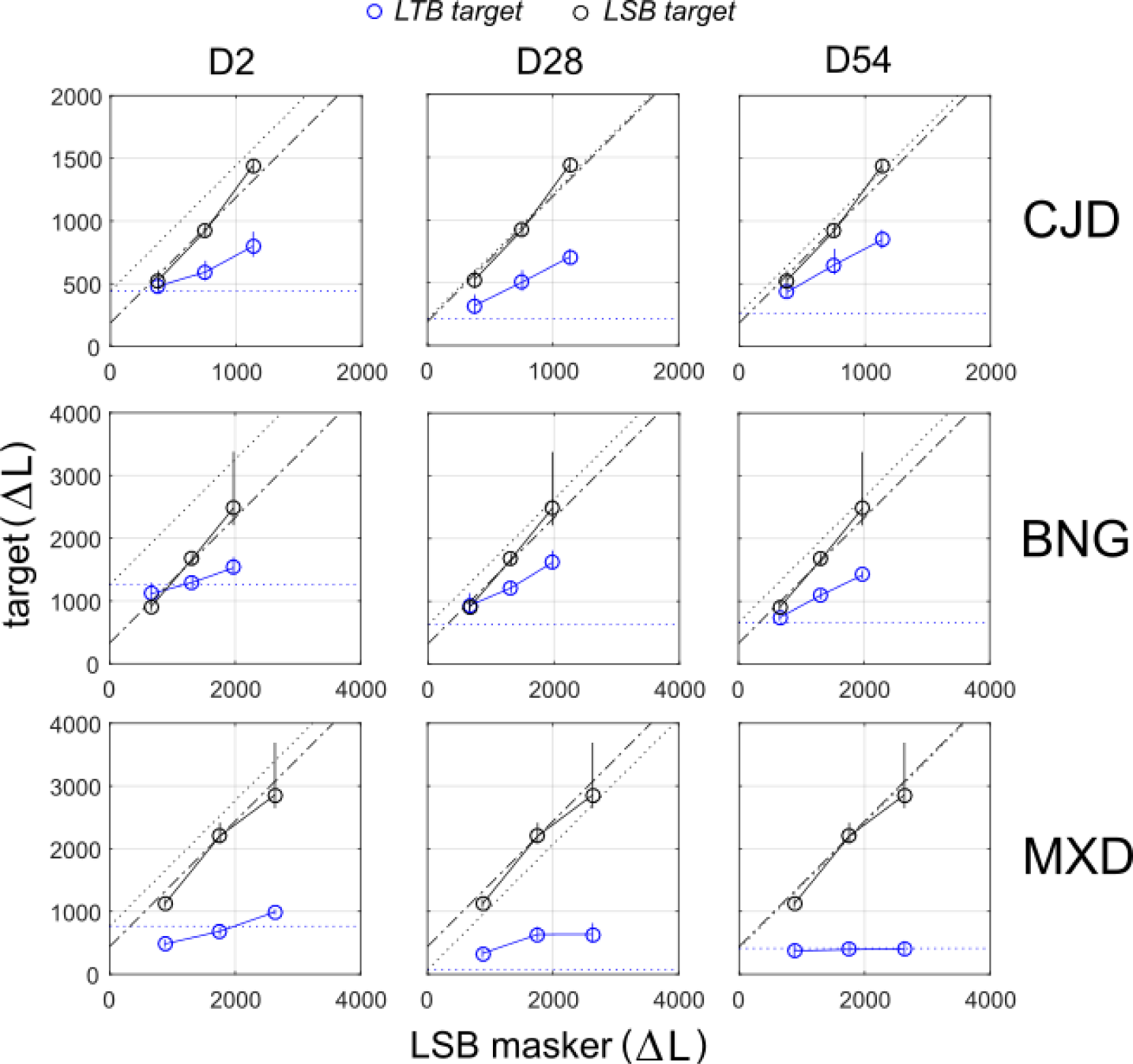
Effects of LSB masker on natural LTB segmentation thresholds. Segmentation thresholds are given in units of absolute luminance differences ΔL across the diagonal for an LTB target (blue symbols) and LSB target (black symbols) masked by an LSB. Blue dotted lines indicate segmentation JND for the LTB target in the absence of a masker (**Experiment 1**). Black dotted lines indicate the predictions of the linear transducer model for the LTB target in the presence of the LSB masker. Black dash-dot lines indicate predictions of the linear transducer model for the LSB target in the presence of the LSB masker. Symbols indicate medians, and lines represent 95% bootstrapped confidence intervals.

Although the LSB may have some masking effect on the LTB, the more interesting question is whether or not the extent of masking is that which we would expect if identical mechanisms were being employed to process both stimuli. We see in **Fig. 6** that the extent to which the LSB masks the LTB is far less than that predicted by a linear transducer model of masking positing identical mechanisms, indicated by the dotted black line. In fact, the observed threshold elevation for LTB segmentation (blue symbols) is significantly less than the model prediction for every observer and for every masker level.

**Fig. 6** also plots the results of **Experiment 3**, and we see that the LSB mask interferes with the segmentation of LSB targets to a much greater extent (black symbols) than LTB targets (blue symbols). In contrast to the LTB targets, the effects of the LSB mask on LSB targets is almost exactly that which would be predicted by the model positing identical mechanisms (black dash-dot line). This is unsurprising of course, since the target and masker are of identical type and will by necessity activate the same underlying mechanisms. From **Experiments 2** and **3**, we can reasonably conclude that there are mechanisms available for LTB segmentation which are relatively unaffected by the presence of the LSB.

By design, this experiment systematically manipulated the relative orientation of the LTB target and LSB masker. **Fig. 7a-c** shows the results of breaking down the trials by the phase relations between the masking LSB and the target LTB, with an organization similar to **Fig. 6**. We see that relative to performance in the incongruent (**inc**) trials, where target and masker have opposite orientations, performance in general is slightly improved (lower thresholds) in the condition where the masker and target are phase-aligned (**con-0,** magenta), and slightly worse (higher thresholds) in the condition where they are opposite-phase (**con-180,** cyan). However, it is important to note that even for the opposite-phase trials, segmentation performance is still markedly better than that which we would predict from our model postulating identical mechanisms (dashed black lines). This relative effects of phase alignment and mis-alignment are somewhat clearer when results are averaged across textures (**Fig. 7b**) or observers (**Fig. 7c**).

**Figure 7:**
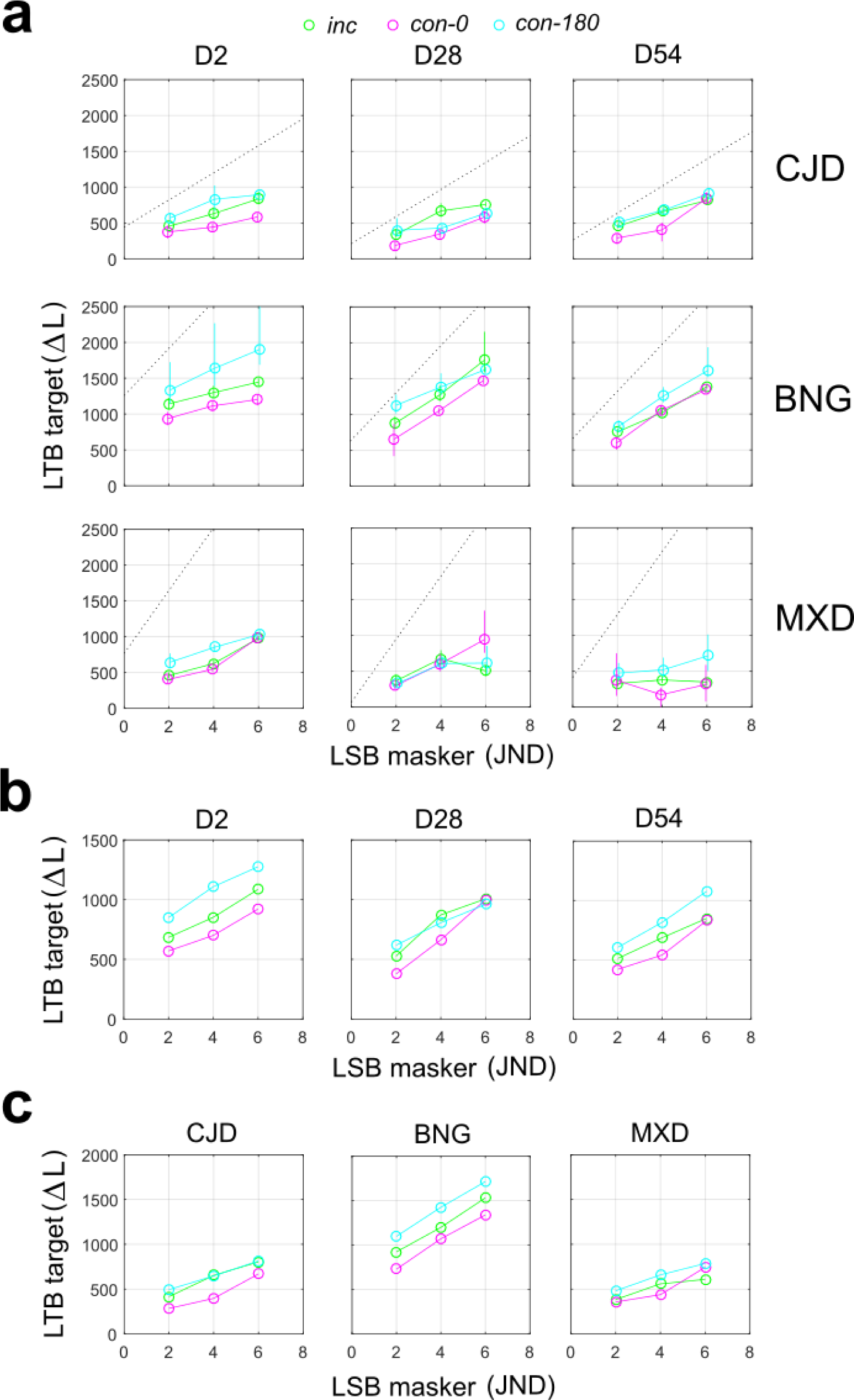
Breakdown of masking results for natural textures by relative orientation and phase of the masker and target. (a) Segmentation thresholds in units of absolute luminance differences ΔL across the diagonal for LTB target with an LSB masker, broken down by the relative orientation and phase of the target and masker (see Fig. 3b for conditions). Here we plot LSB masker level in JND units. Symbols show median thresholds, lines show 68% bootstrapped confidence intervals, and the masker luminance values have been slightly displaced horizontally for visual clarity. Black dotted lines show the predictions of linear transducer model assuming common mechanisms. *Green:* Masker and target have opposite, or incongruent (**inc**) orientations. *Magenta:* Masker and target have same orientation and are phase-aligned (**con-0**). *Cyan:* Masker and target have same orientation and opposite-phase (**con-180**). (b) ) Same as (a), but averaged across observers for each texture. (c) ) Same as (b), but averaged across textures for each observer.

**Fig. 8** shows the difference between the thresholds in the **con-0** and **con-180** cases for all observers and textures, with 95% confidence intervals for the difference. We see that the threshold is generally lower for the same-phase case (**con-0**) than opposite-phase case (**con-180**), with 5/9 cases reaching statistical significance (i.e., 95% confidence interval does not contain zero) for observer CJD, 7/9 for BNG and 3/9 for MXD. In no case was the threshold significantly lower in the **con-180** case than **con-0** for any observer. This is consistent with our previous study, which demonstrated improved segmentation performance for situations where the masking LSB is phase-aligned with the target LTB (**DiMattina & Baker, 2021**). Pooling across 3 textures and 3 masking levels for each observer, we found statistically significant (sign-rank test, N = 9) differences in median segmentation thresholds (**con-0** versus **con-180**) for observers CJD (*p* = 0.004) and BNG (*p* = 0.004), but not MXD (*p* = 0.074).

**Figure 8:**
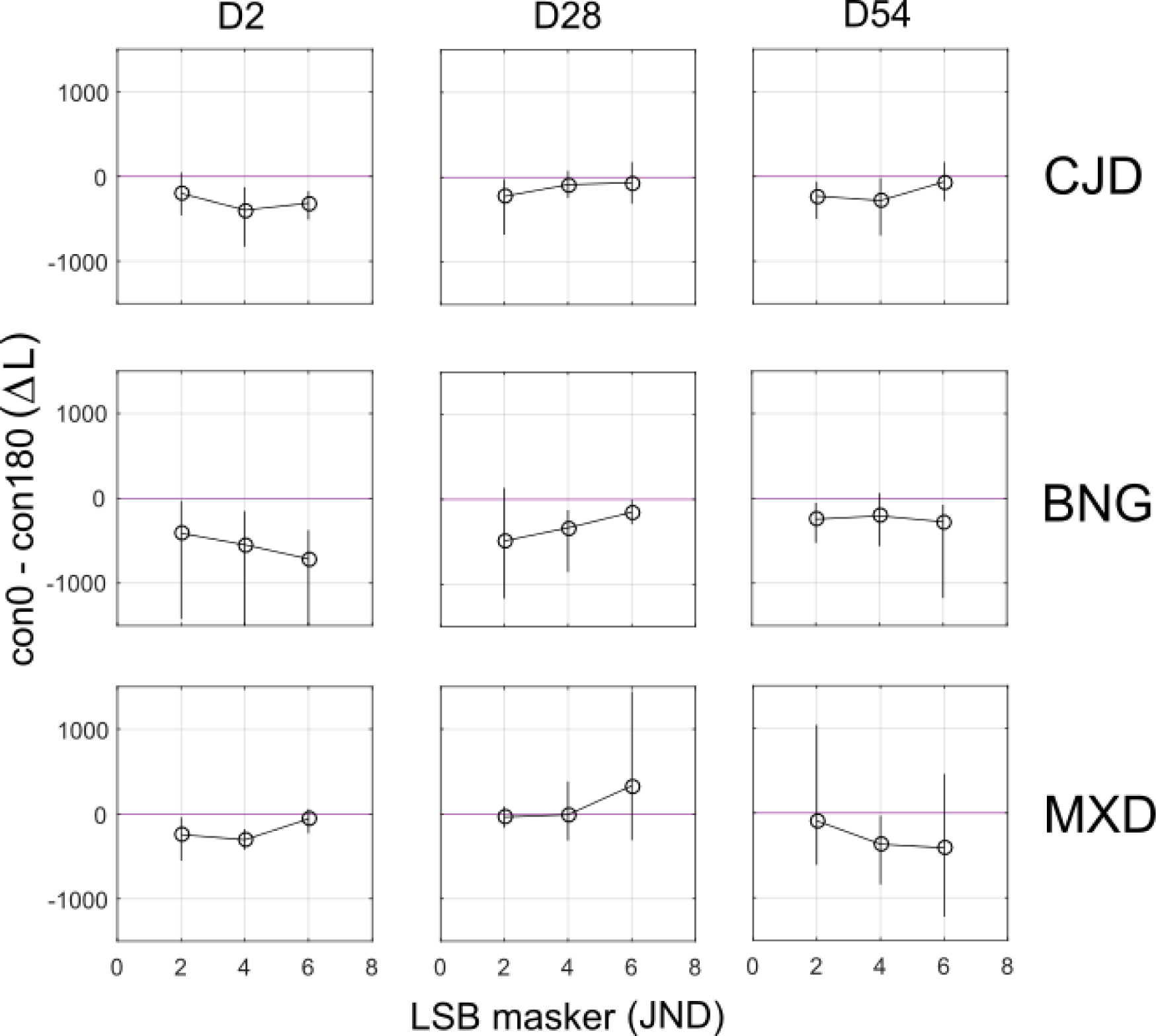
Difference between phase-aligned and opposite-phase masking conditions. Plots show the median difference in the segmentation thresholds between the phase-aligned (**con-0**) and opposite-phase (**con-180**) conditions of **Experiment 2a**. Symbols indicate medians, and lines indicate 95% bootstrapped confidence intervals.

In **Experiment 2a**, we balanced the luminance difference across the anti-diagonal to be zero. We performed an additional control experiment (**Experiment 2b**) in which we do not perform this balancing with one texture (**D54**). Similar results were observed as in **Experiment 2a**, with the curves in the two conditions (blue: **bal**, green: **raw**) exhibiting almost perfect overlap, as shown in **Fig 9a**. Taking the difference in segmentation thresholds between the two cases (**Fig. 9b**), no significant differences were observed for observers CJD or MXD, and for BNG no differences were observed for 2/3 masker levels. More importantly, we see that as in **Experiment 2a** the effects of the masking LSB are far less than predicted by the model assuming a common mechanism (black dotted lines). Consistent with our findings in **Experiment 1**, we conclude that our anti-diagonal luminance balancing procedure has no effects on our main results, and can most likely be omitted in future investigations.

**Figure 9:**
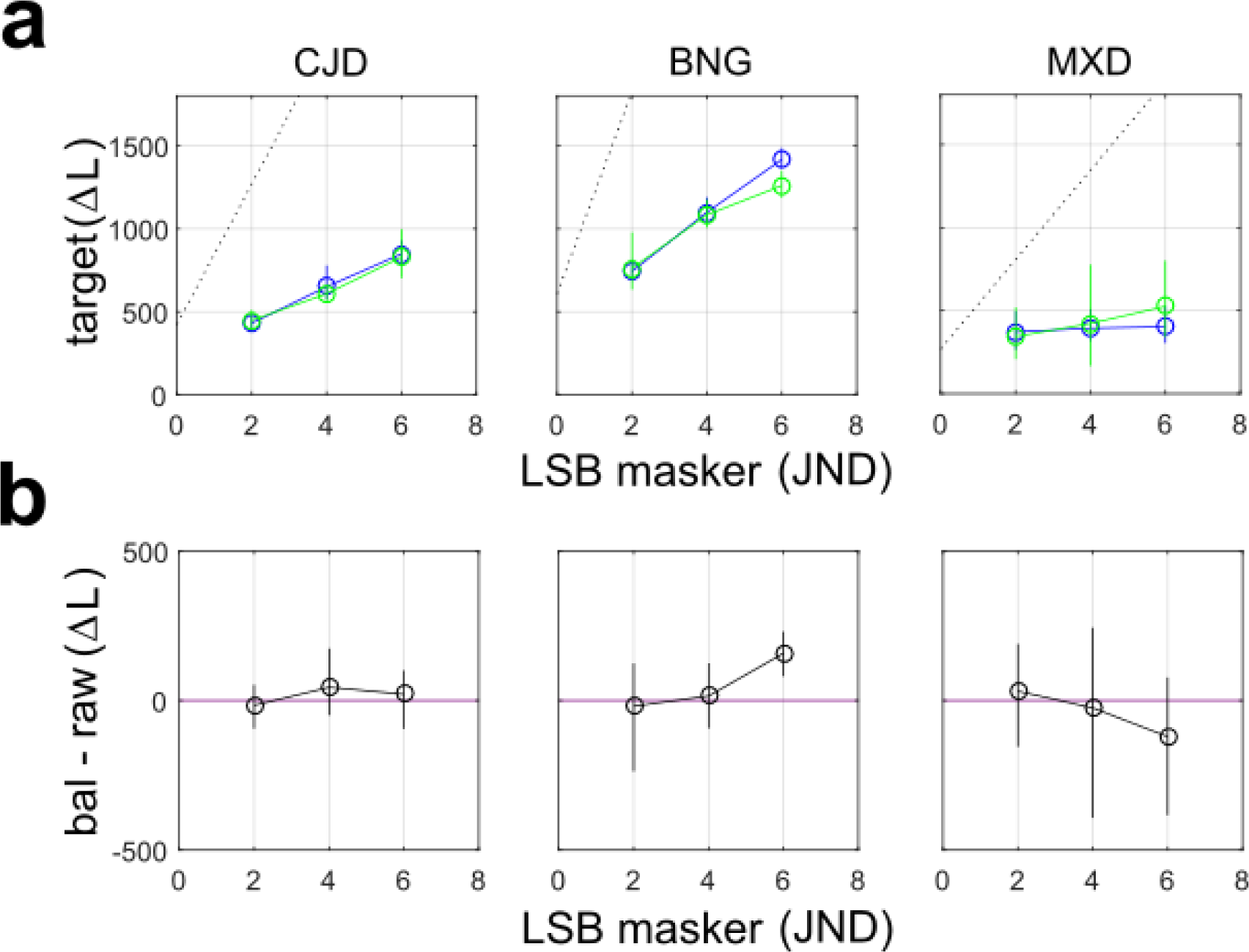
Comparison of LTB segmentation thresholds for texture **D54** in the balanced (**bal**) and unbalanced (**raw**) conditions. (a) Performance in the case where the anti-diagonal is luminance balanced (**bal**) is shown by the green symbols. Performance when the anti-diagonal is not luminance-balanced (**raw**) is shown in blue. Blue and green lines show segmentation thresholds in absence of any masker for both **bal** (blue) and **raw** (green) conditions. Lines show 95% bootstrapped confidence intervals. (b) Difference between the segmentation thresholds in the two cases, plotted in units of absolute luminance difference. Lines show 95% bootstrapped confidence intervals.

### Experiment 4: Masking artificial LTBs with LSBs

Finally, in order to better connect with our previous work (**DiMattina & Baker, 2021**), which utilized artificial LTB stimuli comprised of black and white dots (**Fig. 1b**), we investigated the ability of observers to segment artificial LTB stimuli in the presence of supra-threshold masking LSBs. As we see in **Fig. 10**, our results are largely consistent with those obtained with natural LTB stimuli at both tested densities of Gaussian “dot” micropatterns (*np* = 32, 64 dots per side). The observed thresholds fall well below those predicted by the assumption of identical mechanisms (dashed black lines), consistent with **Experiment 2**.

**Figure 10:**
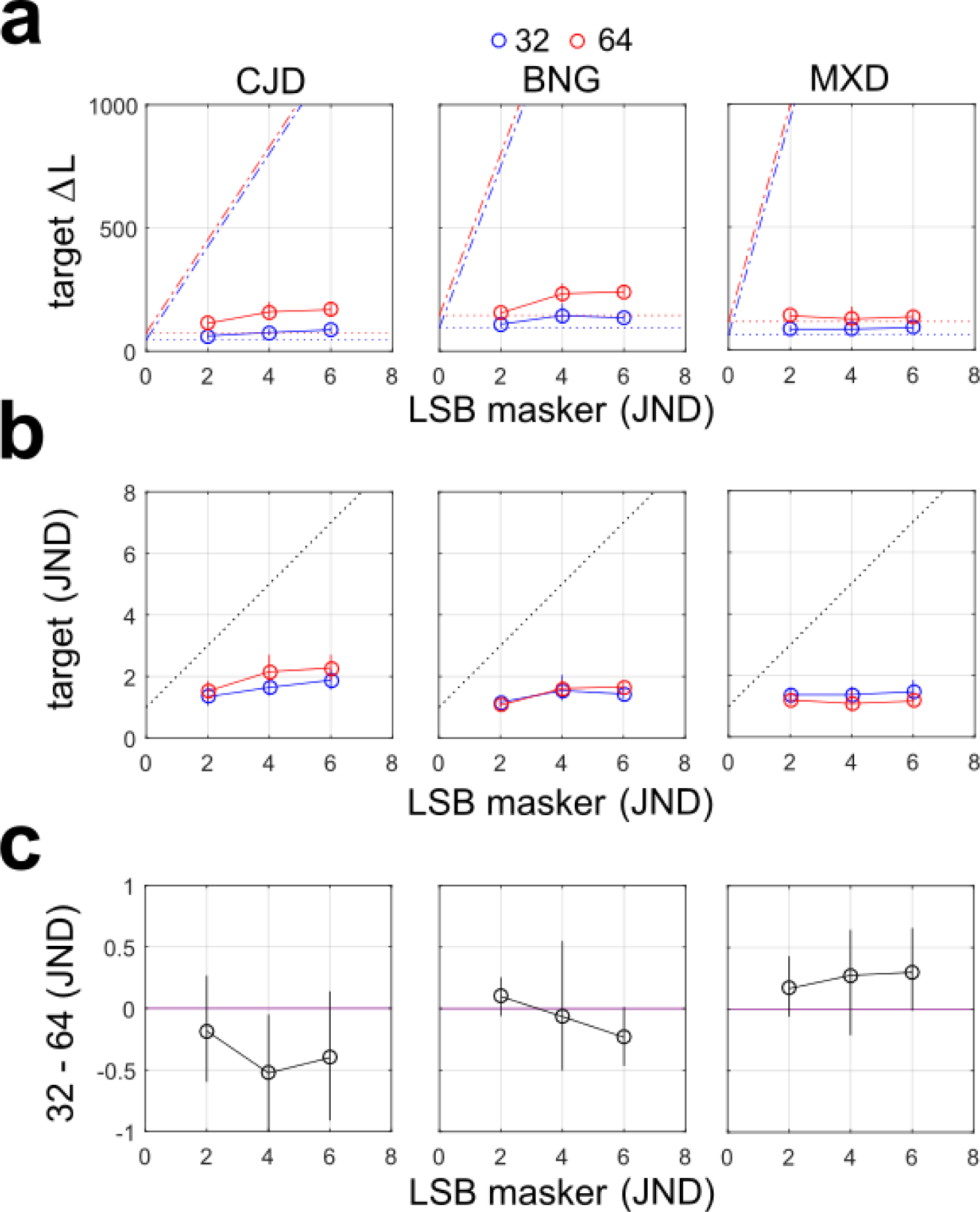
Segmentation thresholds for artificial LTBs masked by LSBs. (a) Segmentation thresholds in units of absolute luminance difference across the diagonal (ΔL) for artificial LTBs (Fig. 1b) with *np* = 32 (blue symbols) and 64 (red symbols) micropatterns per side. Lines show 95% confidence intervals. Dotted lines show segmentation thresholds in the absence of an LSB masker. Dash-dot lines show the predictions of the linear transducer model. (b) ) Same as (a), but in units of JND. (c) Difference between segmentation thresholds for the two densities, in units of JND.

Interestingly, in the absence of any masker, we observed significantly higher segmentation thresholds for the stimuli (in units of ΔL) with 64 micro-patterns (**Fig. 10a**, red dashed lines) than 32 micro-patterns (**Fig. 10a**, blue dashed lines) per side (sign-rank test, *p* < 0.001 for all observers). This difference in segmentation thresholds (in ΔL units) persisted with the maskers present as well (**Fig. 10a**, solid lines). In order to quantify the effect of the maskers relative to the baseline segmentation threshold for each target density, we re-plotted the segmentation thresholds in units of JND with respect to the unmasked case in **Fig. 10b**, and here we see much greater overlap of the curves for the two densities. Taking the difference between the two conditions in JND units (**Fig. 10c**), and pooling across 3 observers and 3 masker levels, for 8/9 cases we see no difference in threshold elevation for the two densities. Therefore, we conclude that the masker has similar effects at the two densities, when normalized by the segmentation JND for that density in the absence of any masker.

Breaking down trials by relative phase/orientation of the LSB masker and LTB target (**Fig. 11a**), we see very similar trends for the different conditions (**inc**, **con-0**, **con-180**) as we see for the natural textures (**Fig. 7**). Although we see a general trend for lower thresholds in the phase-aligned (**con-0**) case than the opposite-phase (**con-180**) case (**Fig. 11b**), for *np* = 32 micropatterns none of these comparisons reach statistical significance (0/9). For *np* = 64 micropatterns only 2/9 comparisons reached significance.

**Figure 11:**
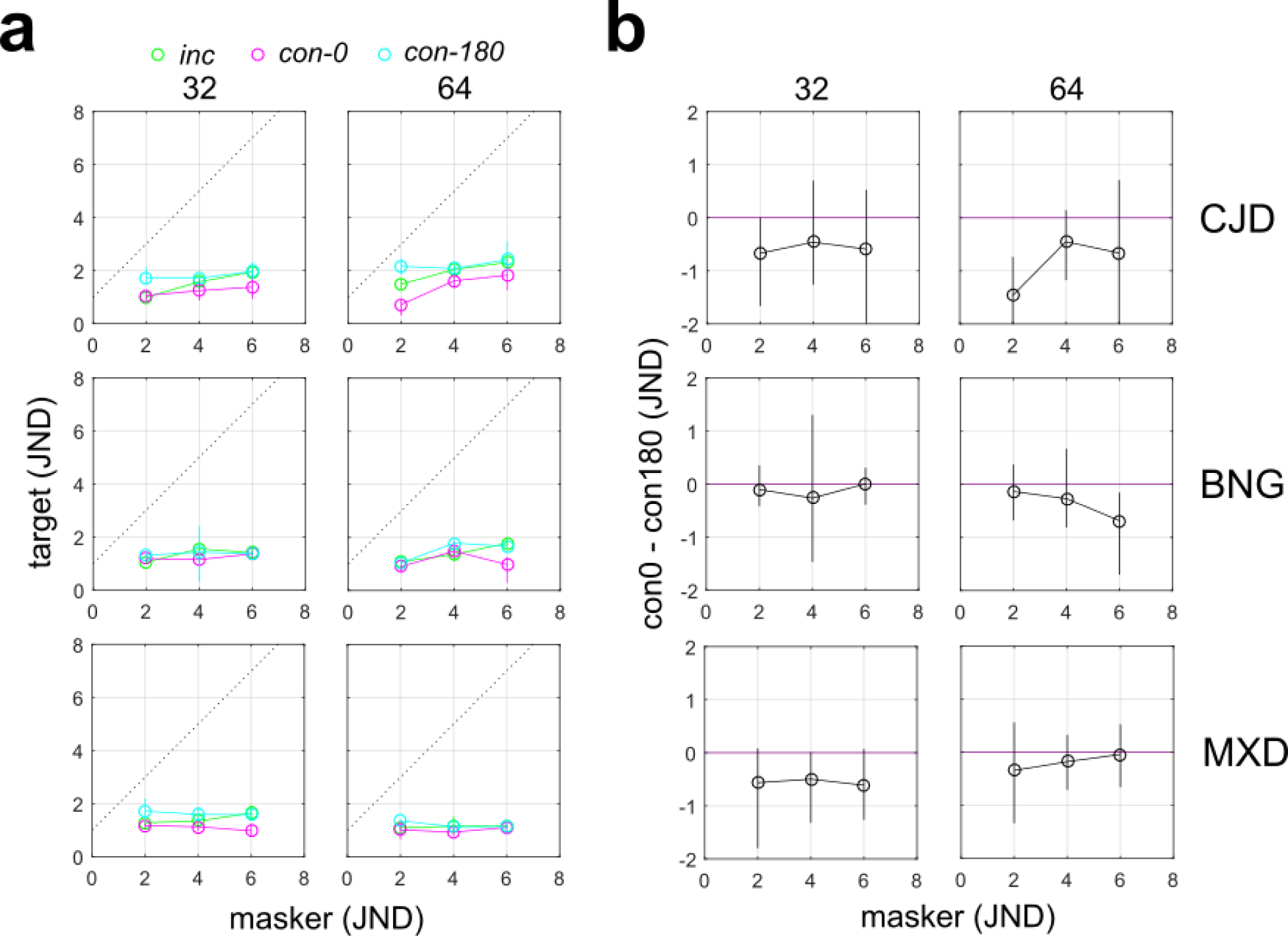
Breakdown of **Experiment 4** by relative phase and orientation for artificial LTB targets **(a)** Same organization as Fig. 7a, but in JND units. **(b)** Same organization as Fig. 8, but in JND units.

## DISCUSSION

### Overview and summary of contributions

In natural images, multiple visual cues for boundary segmentation are available, including texture, color and luminance (**Mely et al., 2016; DiMattina et al., 2012; Martin et al., 2004**). In previous work (**DiMattina & Baker, 2021**), we introduced the conceptual distinction between luminance texture boundaries (LTBs) and luminance step boundaries (LSBs), and examined LTB segmentation in the presence of a masking LSB. We found that observers were robust to interference from LSBs when segmenting LTBs, and that this could be easily explained by positing a two-stage “Filter-Rectify-Filter” (FRF) mechanism for LTB segmentation, similar to models commonly used to explain texture segmentation in the absence of first-order (luminance) cues (e.g., **Malik & Perona, 1990; Landy, 2004; Zavitz & Baker, 2013, 2014; DiMattina & Baker, 2019**). Since one can detect LSBs with only a single stage of linear filtering (**Elder & Sachs, 2004**), our results were consistent with the possibility that these two kinds of boundaries (LTBs/LSBs) may be processed using different (but possibly interacting) underlying neural mechanisms. However, this earlier study suffered from several limitations. In particular, we only examined the robustness of LTB segmentation to interference from masking LSBs when the masker was presented at segmentation threshold. However, in natural vision we typically segment surfaces in the presence of interfering luminance cues arising from clearly visible shadows (**Mammasian, Knill, & Kersten, 1998; Kingdom, Beauce, & Hunter, 2004; Casati & Cavanaugh, 2019**). Furthermore, in **DiMattina & Baker (2021)**, only artificial LTB stimuli like those in **Fig. 1b** were employed, potentially limiting the applicability of our earlier results to understanding natural vision.

In the present study, we adduce further evidence which is consistent with our hypothesis of different mechanisms for LTBs and LSBs. We develop a procedure for taking a uniform natural texture and creating a desired luminance texture difference across a boundary. Creating naturalistic LTBs in this manner from a single texture, rather than simply juxtaposing different textures having different mean luminance (e.g., **Fig. 1a**), helps to minimize the possibility of observers employing non-luminance texture segmentation cues. We compare segmentation thresholds for these natural LTBs to those for these same textures with super-imposed LSBs of the kind that might be caused by a cast shadow. We find across three observers that in the majority of cases, segmentation thresholds for the LTB stimuli are significantly lower than the LSB stimuli, and in no case does the opposite hold. This difference is certainly consistent with the possibility that different mechanisms may be involved in segmenting these kinds of stimuli.

We next performed a series of masking experiments in order to further investigate our hypothesis of multiple mechanisms for luminance boundary cues. We considered the case of a supra-threshold LSB masker which observers were instructed to ignore, and its effects on segmenting both natural (**Experiment 2**) and artificial (**Experiment 4**) target LTB stimuli. We found that the LTB segmentation thresholds were far lower than those which would be predicted by a model assuming a common mechanism for all kinds of luminance differences. By contrast, in a control experiment (**Experiment 3**), we verified that our model does accurately predict performance when the masker and stimulus are of the same kind (LSBs) and necessarily activate identical mechanisms. The results of these masking experiments are also consistent with our previous work (**DiMattina & Baker, 2021**) and support our hypothesis of separate mechanisms for different kinds of luminance boundaries. However, it is important to qualify this by noting that these mechanisms are not entirely independent, as we did observe modest effects of the relative phase of the target and masker, also consistent with our previous study.

### Possible Mechanisms

Our psychophysical results from suggest that the mechanisms of LTB boundary segmentation must be resistant to influence from interfering LSB masker stimuli, even when these maskers give rise to far greater luminance difference than the LTB target. Therefore, we can most likely rule out a simple global luminance difference computation, as such a model would be highly susceptible to interference from masking LSBs (**DiMattina & Baker, 2021**). One possible model consistent with our data is shown in **Fig. 12a**. This model is comprised of two stages of filters: A set of localized, first-stage center-surround filters which detect the individual black and white micro-patterns, followed by a second stage of filtering which integrates broadly over the entire boundary. The first-stage of filtering divides the image (**Fig 12a**, *left*) into separate ON- and OFF- channels, whose outputs are then passed through a half-wave rectifying nonlinearity (purple boxes, *center left*), an essential operation for distinguishing micropatterns having identical shapes but opposite polarities (**Malik & Perona, 1990**). This process yields two new images (*center*) to which one applies large-scale filters defined on the scale of the entire image which are looking for L/R- oblique boundaries in these new images (*center right*). As we see from the responses of these second-stage filters (**Fig. 12a**, *right*), there is a strong response along the left-oblique diagonal, but not along the right-oblique diagonal (responses here correspond to the edge formed by the disc and background grayscale). This becomes more clearly apparent when we pool across the ON- and OFF- channels, as in **Fig. 12b**.

**Figure 12:**
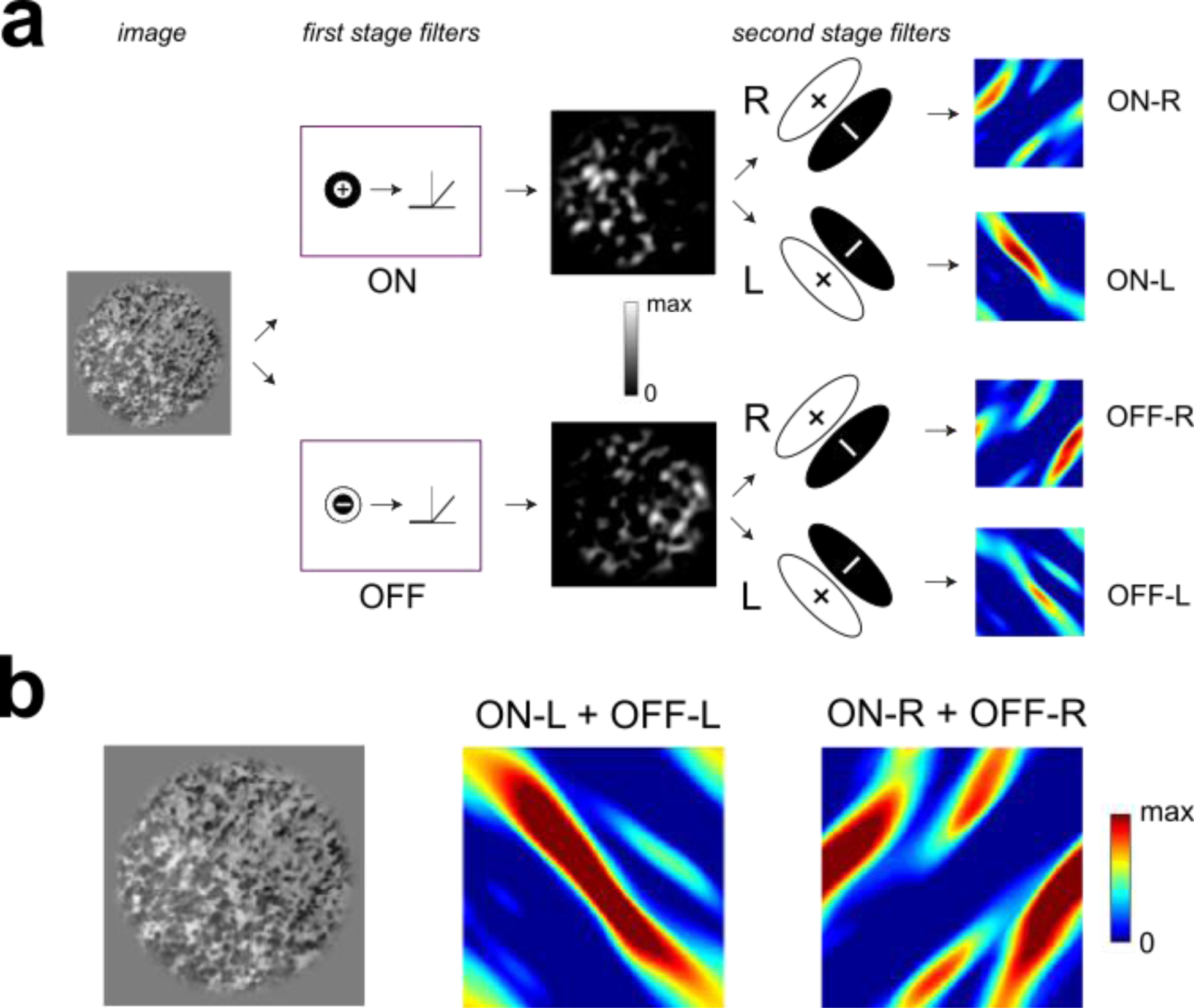
Hypothetical model with two filtering stages which is sensitive to LTBs and not sensitive to LSBs. The input image to this model has an LTB oriented left-oblique and a super- imposed LSB which id oriented right-oblique, with both diagonals having a luminance difference of ΔL = 1000. (a) Schematic of the model. The input image (*far left*) is convolved with ON-center and OFF-center first-stage filters defined at a small spatial scale (*center-left*). The half-wave rectified output of this stage is plotted in grayscale (*center*), and we see here that the ON-channel responds more strongly to the lower left, and the OFF-channel more to the upper right. The outputs of each channel are then passed to large scale second-stage filters defined on the scale of the whole edge (*center-right*). These second-stage filters look for L/R-oblique boundaries, and here we see (*far right*) that the filters sensitive to L-oblique boundaries show a strong response along the diagonal (scale bar in (b)). (b) Combined responses (ON + OFF) of the second-stage filters looking for left-oblique (*center*) and right-oblique (*right*) boundaries. Original image shown on *left*.

Assuming that the first-stage filters respond weakly to uniform luminance, a model like that in **Fig. 12a** would be able to detect luminance texture boundaries with minimal interference from luminance steps. However, if the first-stage filters exhibit a non-zero DC response, there is the potential for the phase of a masking LSB having a congruent boundary orientation to influence segmentation thresholds, as we observed in our data (**Fig. 7**) and in our previous work (**DiMattina & Baker, 2021**). However, this is by no means the only possible explanation for our observed interactions between LSBs and LTBs. It may also be the case that the first-stage filters are DC balanced and insensitive to luminance steps, and LTB and LSB information is first combined at a later stage of processing. It remains for future work to develop experiments to distinguish between these interesting possibilities.

## Limitations and Future Directions

Edges in luminance can arise from many causes in natural scenes, including surface changes (i.e., occlusions), material changes, surface orientation changes, and cast shadows (**Vilankar et al., 2014**). In this work, we have suggested that there may be different mechanisms for LTB and LSB segmentation, and these different mechanisms may have functional implications for distinguishing luminance edges arising from cast shadows from those arising from surface boundaries. However, in order to test this explicitly, one would have to implement these different hypothesized biological mechanisms using computer simulations, and test the ability of each mechanisms to distinguish edges in natural images arising from shadows from edges arising from natural surface boundaries. Ideally, the performance of these operators should be compared to human psychophysical performance on the task of distinguishing shadows from surface boundaries, as in previous studies (**Breuil et al., 2019**). To the best of our knowledge, such a computational study testing biologically plausible mechanisms on this task has yet to be conducted, and we are currently performing computational studies to evaluate the ability models like that shown in **Fig. 12a** to distinguish edges arising from occlusion boundaries and cast shadows in grayscale images.

Another limitation of the current work is that in natural vision, there are a number of cues which can potentially distinguish shadows from surface boundaries, including orientation (**Wolfson & Landy, 1998**), micro-pattern density (**Zavitz & Baker, 2014**), and contrast (**Dakin & Mareschal, 2000**; **DiMattina & Baker, 2019**). It is of interest for future work to better understand how luminance cues (LTBs and LSBs alike) interact with these various second-order texture cues for boundary segmentation. This can be accomplished using various sub-threshold summation paradigms and fitting quantitative models of cue combination to the data (e.g., **Motoyoshi & Nishida, 2004; Kingdom et al. 2015**), and/or using masking paradigms like those considered here in which observers must segment using one cue while ignoring the other (**Saarela & Landy, 2012**). Previous psychophysical work studying the interaction of first and second-order cues has focused on detection tasks, and these results have suggested separate mechanisms for processing first- and second-order cues (**Schofield & Georgeson, 1999; Allard & Faubert, 2007**). By contrast, neurophysiological investigations have suggested the existence of “form-cue invariant” responses which respond to boundaries defined by either first or second-order cues (**Li et al, 2014**), and even demonstrated nonlinear interactions between these cues (**Hutchinson, Ledgeway, & Baker, 2016**). Of particular interest for both psychophysics and physiology is comparing the interactions between first-order and second-order information in two cases: (1) Where the first-order cue is defined by texture (LTB), as might be the case for two different surfaces, and (2) The first-order cue is a luminance step (LSB), which can possibly be caused by a shadow. One reasonable hypothesis is that we may observe a greater degree of summation between first- and second-order cues in case (1) than in case (2), since changes in texture micro-pattern properties provide stronger evidence for a surface boundary. Similarly, we may expect a greater degree of masking or interference between first- and second-order cues in case (1) than in case (2) as well. These intriguing possibilities remain to be investigated in future experimental work.

Surface boundaries in natural scenes are generally accompanied by changes in luminance, chromatic, and texture cues (**Mely et al., 2016; Hansen & Gegenfurtner, 2009; DiMattina et al., 2012; Vilankar et al., 2014**). By contrast, shadows only give rise to changes in luminance, and sometimes these luminance changes can be gradual due to penumbral blur (**Casati & Cavanagh, 2019**). Therefore, mechanisms sensitive to changes in chromaticity and/or texture can potentially contribute to distinguishing the cause of an edge. Indeed, previous work has demonstrated that color information can help observers determine whether or not an edge is caused by a shadow or change in material (**Breuil et al., 2019**), or determine whether two regions are from the same surface (**Ing et al., 2010**). Likewise, when texture information is removed from a natural occlusion boundary, it becomes harder to accurately identify as a boundary (**McDermott, 2004; DiMattina et al., 2012**). The bulk of studies investigating texture segmentation have only considered textures defined by “second-order” cues, with luminance differences held constant. However, patterns of luminance are an important component of texture and can even serve to define textures in the absence of other cues (**Fig. 1b**). We hope that the present paper and related work (**DiMattina & Baker, 2021**) represents a modest first step in the direction of a better and more complete understanding of the role of luminance differences in texture boundary segmentation.

## Acknowledgements

We thank FGCU undergraduates in the Computational Perception Lab for help with data collection.

